# Mutant EGFR is a preferred SMURF2 substrate of ubiquitination: role in enhanced receptor stability and TKI sensitivity

**DOI:** 10.1101/2020.04.02.022012

**Authors:** Paramita Ray, Krishnan Raghunathan, Aarif Ahsan, Uday Sankar Allam, Shirish Shukla, Venkatesha Basrur, Sarah Veatch, Theodore S. Lawrence, Mukesh K. Nyati, Dipankar Ray

**Affiliations:** Departments of Radiation Oncology, The University of Michigan Medical School, Ann Arbor, Michigan 48109; Biophysics, The University of Michigan Medical School, Ann Arbor, Michigan 48109; Pathology, The University of Michigan Medical School, Ann Arbor, Michigan 48109

**Keywords:** Epidermal growth factor receptor (EGFR), Smad ubiquitination regulatory factor 2 (SMURF2), Ubiquitin conjugating enzyme H5 (UBCH5), protective ubiquitination, tyrosine kinase inhibitor (TKI) resistance

## Abstract

We previously reported that differential protein degradation of TKI-sensitive [L858R, del(E746-A750)] and resistant (T790M) epidermal growth factor receptor (EGFR) mutants upon erlotinib treatment correlates with drug sensitivity. However, the molecular mechanism remains unclear. We also reported SMAD ubiquitination regulatory factor 2 (SMURF2) ligase activity is important in stabilizing EGFR. Here, using *in vitro* and *in vivo* ubiquitination assays, mass spectrometry, and super-resolution microscopy, we show SMURF2-EGFR functional interaction is critical in receptor stability and TKI sensitivity. We found that L858R/T790M EGFR is a preferred substrate of SMURF2-UBCH5 (an E3-E2) complex-mediated K63-linked polyubiquitination, which preferentially stabilizes mutant receptor. We identified four lysine (K) residues (K721, 846, 1037 and 1164) as the sites of ubiquitination and replacement of K to acetylation-mimicking asparagine (Q) at K1037 position in L858R/T790M background converts the stable protein sensitive to erlotinib-induced degradation. Using STochastic Optical Reconstruction Microscopy (STORM) imaging, we show that SMURF2 presence allows longer membrane retention of activated EGFR upon EGF treatment, whereas, siRNA-mediated *SMURF2* knockdown fastens receptor endocytosis and lysosome enrichment. In an erlotinib-sensitive PC9 cells, SMURF2 overexpression increased EGFR levels with improved erlotinib tolerance, whereas, *SMURF2* knockdown decreased EGFR steady state levels in NCI-H1975 and PC9-AR cells to overcome erlotinib and AZD-9291 resistance respectively. Additionally, by genetically altering the SMURF2-UBCH5 complex formation destabilized EGFR. Together, we propose that SMURF2-mediated preferential polyubiquitination of L858R/T790M EGFR may be competing with acetylation-mediated receptor internalization to provide enhanced receptor stability and that disruption of the E3-E2 complex may be an attractive alternate to overcome TKI resistance.

## INTRODUCTION

There are two “classical” activating mutations, in-frame deletions in exon 19 and a point mutation in exon 21 (L858R)] in *EGFR*, which drive adenocarcinoma of the lung in the majority of never smokers (1). The presence of such receptor mutations is a marker for sensitivity to treatment with EGFR tyrosine kinase inhibitors (TKIs). First generation TKIs, such as erlotinib and gefitinib, produce responses, however, within a year, tumors develop TKI resistance. In about 50% of these cases, resistance is due to the enrichment of a second point mutation at exon 20, where threonine at 790 position is mutated to methionine (T790M) (2). To circumvent resistance, a third generation EGFR inhibitor (AZD9291, osimertinib) has been generated, which is effective for patients carrying the T790M mutation (3). Unfortunately, with time, patients show osimertinib resistance due to multiple mechanisms (4). In spite of significant efforts, the molecular mechanisms causing acquired TKI-resistance are still poorly understood. In initial studies, differential drug-binding abilities and altered ATP-binding affinities of various EGFR mutants have been proposed as responsible for differential TKI response (5). In the case of osimertinib resistance, an additional mutation at the tyrosine kinase domain of EGFR (C797S) has been implicated in about one third of cases (6–8), and MET amplification has also been noted (4). However, these factors together may be only a part of the acquired resistance story.

TKI resistant cell lines are sensitive to EGFR degradation or knockdown by RNA interference, which suggests that the physical presence of EGFR independent of its kinase activity is important in cancer cell survival (9,10). These findings led us to hypothesize that inducing EGFR degradation may overcome TKI resistance. We previously reported that EGFR degradation determines the response to radiation and chemotherapy and that targeting EGFR for degradation can induce cytotoxicity in cancer cells (11–18). We further elucidated the importance of EGFR degradation in head and neck cancer patients treated with erlotinib (19). Together these findings led us to hypothesize that EGFR degradation is a key phenomenon in determining cytotoxicity to various chemotherapies, radiation or TKI targeting EGFR. Differential EGFR degradation may contribute to the dramatic difference in sensitivity to TKIs of patients harboring L858R compared to T790M. Our data demonstrated that TKI sensitive EGFR delE746-A759 or L858R mutant proteins undergo rapid degradation upon erlotinib treatment, whereas, TKI-resistant L858R/T790M protein is highly stable (20).

Ubiquitin ligases and deubiquitinating enzymes, which play an important role in EGFR endocytosis, trafficking, and degradation, regulate EGFR stability (21). The most extensively studied E3 ligase responsible for EGF-mediated EGFR polyubiquitination is c-CBL (22). Additionally, AIP4/ITCH, pVHL, and UBE4B play roles in the polyubiquitination and degradation of EGFR (23,24). We have reported that the HECT-type E3 ubiquitin ligase, Smad ubiquitination regulatory factor 2 (SMURF2) directly interacts with EGFR, but unlike several others, SMURF2’s ubiquitin ligase activity is critical for stabilizing EGFR (25). Here, we study the mechanistic and functional importance of SMURF2-mediated ubiquitination on different EGFR mutants, identify residue(s) undergoing post-translational modifications, and determine the relevance of mutant receptor membrane retention and protein stability, which together can predict TKI response. Additionally, we explore the impact of disruption of SMURF2 interaction with its cognate ubiquitin conjugating enzyme (E2), UBCH5 on EGFR protein stability to overcome TKI resistance.

## RESULTS

### EGFR (L858R+T790M) is a preferred substrate for SMURF2-UBCH5 mediated ubiquitination

We previously reported that SMURF2 ubiquitin ligase activity is critical in maintaining EGFR protein stability (25). Mutant EGFR is a preferred SMURF2 substrate of ubiquitination: role in enhanced receptor stability and TKI sensitivity To determine whether the SMURF2 can directly ubiquitinate EGFR and whether such modification may vary depending on the type of EGFR mutations, we performed *in vitro* ubiquitination assay by incubating either wild type (WT) or different EGFR [either L858R (L) or L858R/T790M (L+T)] mutants with SMURF2. As SMURF2 partners with either UBCH7 or UBCH5 as E2, we tested SMURF2 catalytic activity on EGFR in the presence or absence of either one of the E2s. As shown in **Fig. 1A**, we noted substantial polyubiquitination of L+T mutant EGFR compared to WT and L mutant. Additionally, the polyubiquitinated species formed in the presence of UBCH5 was of higher molecular weight compared to those formed in the presence of UBCH7; the later we speculate to be autoubiquitinated SMURF2, as reported earlier (26). To decipher whether preferential binding may be responsible for SMURF2-mediated mutant EGFR ubiquitination, we performed immunoprecipitation studies. As shown in **Fig. 1B**, we noted comparable binding of SMURF2 with WT, L and L+T mutant EGFR. To understand the kind of ubiquitin linkage formed on L+T mutant EGFR, we utilized different ubiquitin mutants (K11R, K48R and K63R) deficient in forming specific linkages. As shown in **Fig. 1C**, there was a significant reduction of mutant EGFR polyubiquitination only in the presence of K63R recombinant ubiquitin, suggesting SMURF2-mediated EGFR ubiquitination is primarily K63-linked.

**Figure 1.**
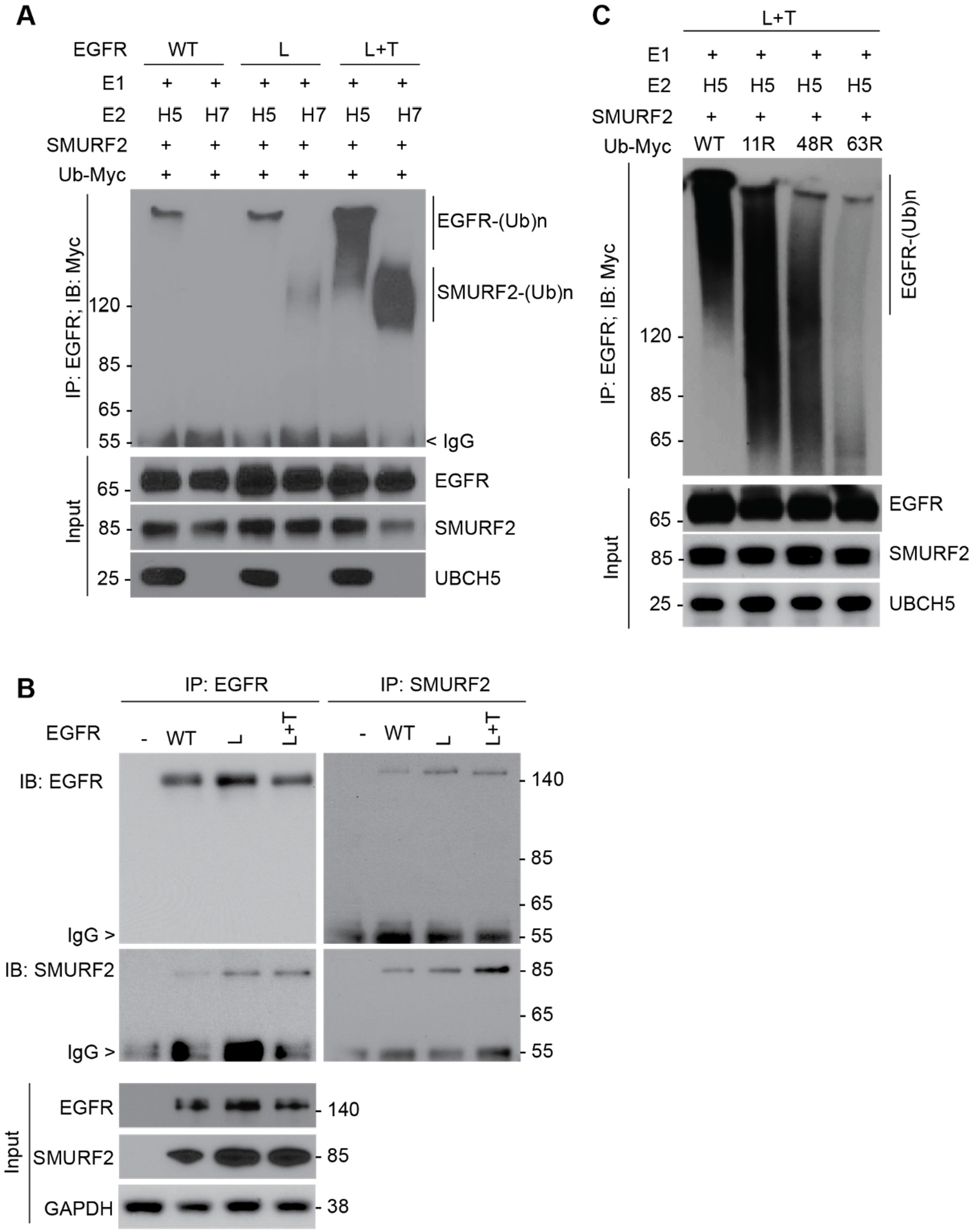
EGFR (L858R/T790M) is a preferred substrate for SMURF2-UBCH5 mediated ubiquitination. **(A)** Purified EGFR proteins, either wild type (WT) or mutants [L858R (L) and L858R/T790M (L+T)] were subjected to *in vitro* ubiquitination using recombinant SMURF2 as an E3 in the presence of either UBCH5 or UBCH7 as E2 enzymes. Following completion, reaction mixtures were subjected to immunoprecipitation using the EGFR antibody followed by immunoblotting using indicated antibodies. **(B)** CHO cells overexpressing EGFR (WT, L and L+T) and SMURF2 were immunoprecipitated with either EGFR or SMURF2 antibodies and immunoblotted for indicated antibodies. **(C)** SMURF2:UBCH5 mediated *in vitro* ubiquitination of mutant EGFR (L+T) was performed as described above in the presence of WT or different ubiquitin mutants (K11R, K48R and K63R) deficient in promoting specific linkages. Higher molecular weight ubiquitinated EGFR species were detected following immunoprecipitation and immunoblotting.

### Acetylation mimicking K1037Q mutation in L+T mutant EGFR background converts the stable receptor vulnerable to TKI-mediated degradation

To determine the site(s) ubiquitinated in EGFR by SMURF2, we performed mass spectrometry analysis of immunoprecipitated receptor and identified four lysine residues K721, K846, K1037 and K1164 (**Supporting Fig. 1A, B, C, D**). Interestingly, K721 and K846 are known to be important for EGFR kinase activity (27–29) and the latter two residues (K1037 and 1164) undergo acetylation necessary for receptor internalization (30), thus indirectly impacting receptor degradation. Acetylation and ubiquitination counteract each other (31). To better understand the role of the two post-translational processes in determining EGFR protein stability, we generated sitespecific EGFR mutants of K1037 and K1164, mutated either to Arginine (R, can be neither ubiquitinated nor acetylated) or Glutamine (Q, mimics constitutive acetylation). As shown in **Fig. 2A**, the incorporation of K to Q mutation in L+T background converted a stable double mutant to unstable following erlotinib treatment with consequent accumulation of polyubiquitinated species (**Fig. 2B**). In contrast, conversion of K to R minimally impacted the triple mutant EGFR steady state levels following erlotinib treatment (**Fig. 2A**).

**Figure 2.**
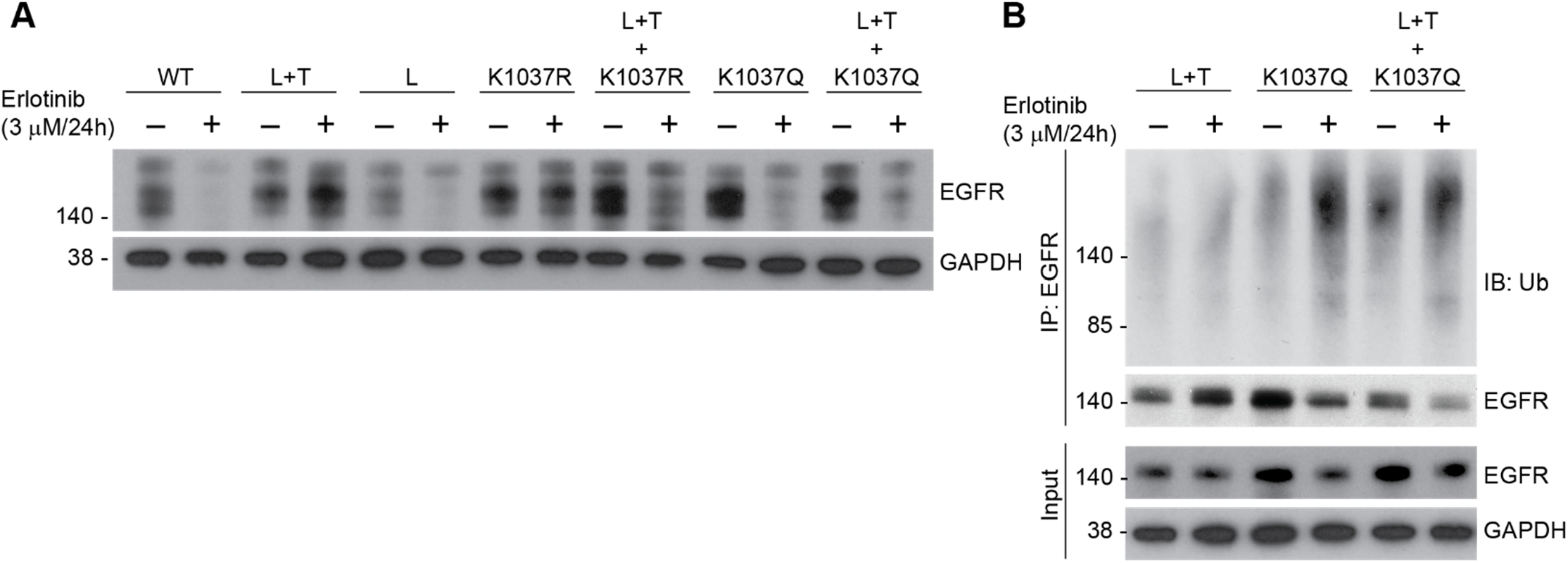
Incorporation of K1037Q mutation in L+T EGFR background converts stable receptor vulnerable to TKI-mediated degradation. **(A)** CHO cells overexpressing either wild type or different mutants of EGFR (as indicated in the figure), were treated with 3 μM erlotinib for 24h. Cell lysates were then subjected to immunoblot analysis with indicated antibodies. **(B)** CHO cells transfected with Indicated EGFR mutants were treated with erlotinib as above followed by immunoprecipitation using EGFR antibody and immunoblotting with indicated antibodies.

### EGFR surface density is dependent on SMURF2 expression

Ubiquitination and acetylation counteract each other and acetylation of EGFR has been associated with receptor internalization. Therefore, we tested the hypothesis that SMURF2-mediated ubiquitination is important to maintain the surface density of EGFR by enhancing protein stability and thereby reducing internalization. We addressed this question by directly measuring the protein density in the membrane using STochastic Optical Reconstruction Microscopy (STORM) (32,33). STORM images were quantified using pair-correlation based analytical method that can correct quantitative artifacts arising from over-counting. This method takes advantage of the observation that the magnitude of self-clustering that arises from over-counting is inversely proportional to the density of the labeled protein, as long as the labeled protein is sampled randomly.

We first tested the validity of our approach by quantifying EGFR levels in the membranes of multiple cell lines (CHO with no detectable EGFR, and multiple head and neck cancer cell lines, UMSCC-11b, UMSCC-1 and UMSCC-29b) those express varied amounts of receptor using conventional immunoblotting (**Supporting Fig. 2A**). As shown (**Supporting Fig. 2B and 2C**), there was a high correlation (R^2^=0.9854) observed between the conventional immunoblotting and Total Internal Reflection Fluorescence Microscopy (TIRFM), demonstrating the applicability. To further validate the methodology, we measured the receptor population in a human breast cancer MCF7 cell line with known (1000-5000 per cell) numbers of EGFR (34). As shown in **Supporting Fig. 2D and 2E**, we counted n=900-1771 receptor molecules per cell (assuming a radius of 20-25 μm) consistent with the literature.

Having established the reliability of the methodology, we next quantified EGFR surface density in the presence and absence of SMURF2. We conducted this experiment both in the presence and absence of EGF. EGF treatment caused a decrease in surface EGFR population arising from internalization of EGFR (**Supporting Fig. 2F**). The decrease in EGFR surface expression was greater following EGF treatment in *SMURF2* siRNA treated cells (~60% vs. ~40% loss) compared to control siRNA treatment, indicating a role for SMURF2 in maintaining EGFR membrane levels.

### EGF treatment promotes EGFR-SMURF2 membrane co-clustering

We previously reported a dynamic interaction between EGFR and SMURF2 (25) and here we found that knockdown of SMURF2 alters EGFR density, especially following EGF stimulation. This led us to hypothesize that EGFR-SMURF2 interaction occurs proximal to the plasma membrane. To test this hypothesis, we conducted immunoprecipitation and immunoblot analysis in cell-fractionated isolates from UMSCC-1 cells, using siRNA to alter SMURF2 levels. As expected, we observed EGFR was predominantly present in the plasma membrane, whereas, SMURF2 was primarily cytosolic (**Fig. 3A**, right panel) with a small fraction present in the membrane. Interestingly, we found an increased EGFR-SMURF2 interaction in the membrane fraction following EGF treatment (**Fig. 3A**, left panel). Next, we conducted two color STORM experiments to quantify interaction between EGFR and SMURF2 localized proximal to the plasma membrane (**Fig. 3B**). UMSCC-1 cells were grown normally in serum containing media and SMURF2 and EGFR levels were determined in the presence and absence of EGF (10ng/ml, 30 min). Interestingly, upon stimulation with EGF, we observed an increase (2.9±0.3 fold) in SMURF2 co-clustering with EGFR to the plasma membrane (**Fig. 3C**). SMURF2 density in the membrane went up almost 4 fold in the membrane in line with the cell fractionation studies.

**Figure 3.**
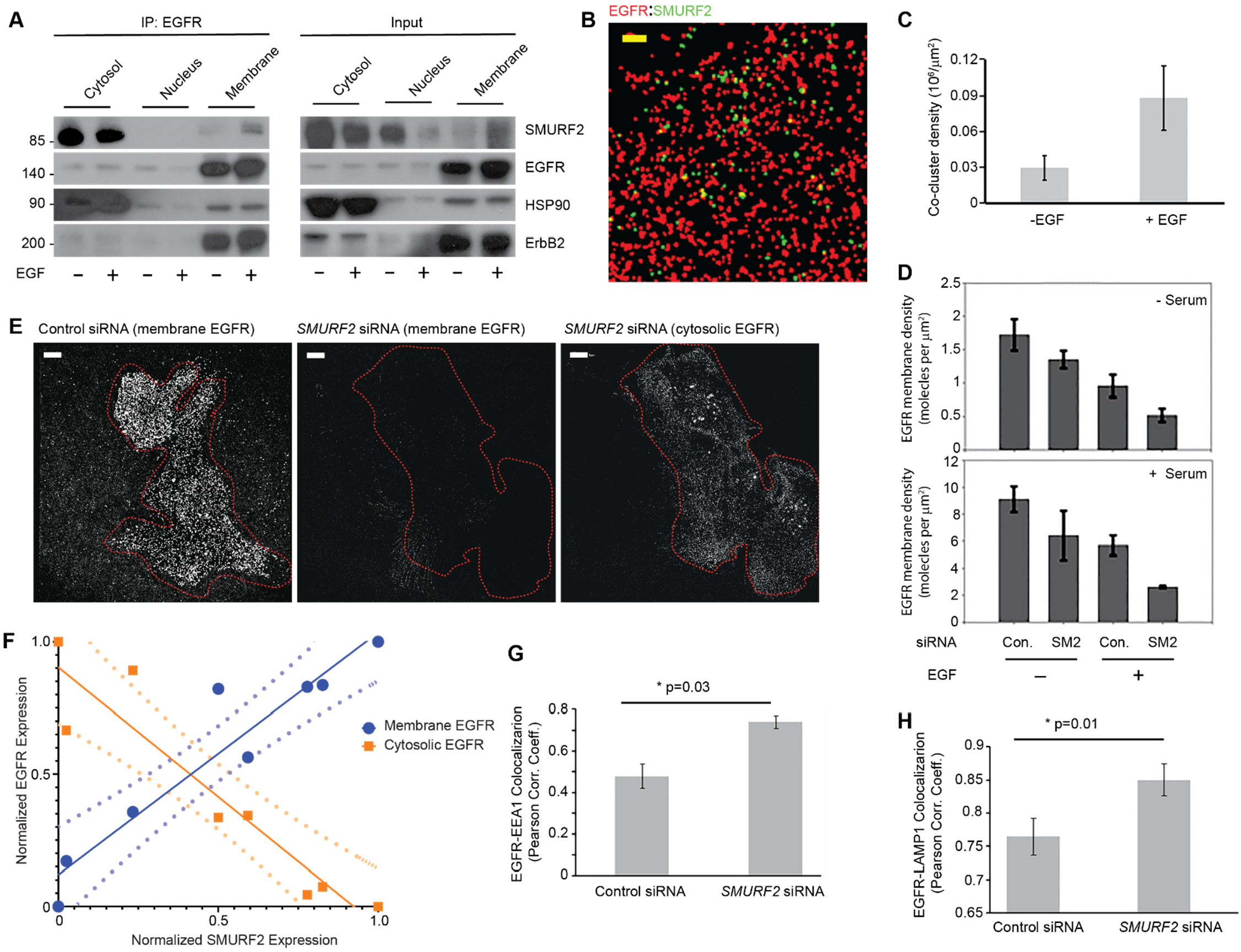
EGFR membrane density is dependent on SMURF2 expression and EGF treatment promotes EGFR-SMURF2 co-clustering. **(A)** UMSCC-1 cells were either left untreated or treated with EGF (10 ng/ml for 6h). Cell lysates were then subjected to fractionation into cytosol, nuclear and membrane fractions. Following, the fractionated samples were immunoprecipitated using EGFR antibody followed by immunoblotting as indicated. **(B)** Representative STORM image showing EGFR and SMURF2 co-clustering on UMSCC-1 cell surface (-EGF; +serum) following immunostaining. Scale bar, 5 μm. **(C)** Increase in EGFR-SMURF2 co-cluster density was observed following EGF treatment (10 ng/ml for 6h). **(D)** Quantification of EGFR membrane density in UMSCC-1 cells transfected either with control or *SMURF2* siRNA and grown in the presence and absence of serum and treated with EGF (10 ng/ml for 30 min) where indicated. **(E)** Membrane-proximal EGFR levels were compared for UMSCC-1 cells transfected with either control or *SMURF2* siRNA using immunofluorescence staining and super-resolution microscopy. Reconstructed image showing EGFR density in plasma membrane of a cell transfected with control siRNA. Reconstructed images of both plasma membrane and cytosolic EGFR of a cell transfected with *SMURF2* siRNA. **(F)** Correlation of membrane and cytosolic EGFR levels with the cytosolic expression of SMURF2 in cells. The relative expression of proteins in cells were quantified using epifluorescence images for SMURF2 and EGFR (cytosol) and by imaging the same cell in the TIRF mode for the membrane expression of EGFR. The dotted lines represent the 95% confidence bound when fitted with a linear regression model. **(G and H)** UMSCC-1 cells transfected with either control or SMURF2 siRNA were co-stained either with EGFR and EEA1 **(in F)** or with EGFR and LASMMP1 **(in G).** Following imaging, Pearson correlation coefficient (PCC) were determined for either EGFR and EEA1 (*p*=0.03) or EGFR (-and LAMP1 (*p*=0.01) as a measure of colocalization between EGFR and the marker. PCC was calculated by drawing a mask outside of the cell and the two conditions (control vs SMURF2 siRNA) were compared for at least in 10 cells for the analysis.

To better understand the protective role of SMURF2 in the retention of activated receptor in the cell membrane, we determined changes in EGFR density in the plasma membrane following *SMURF2* knockdown both in the presence and absence of EGF. Surface EGFR molecules of UMSCC-1 cells grown in serum containing medium were 5.3±0.6 fold higher compared to cells grown in serum-free condition (**Fig. 3D**). As previously noted, EGF (10ng/ml for 30 min) treatment caused a decrease of 44.5±3.3 and 37.2±1.2% respectively for cells grown in serum starved and serum containing media, suggesting that EGF-induced EGFR internalization is independent of the growth conditions. However, the impact of *SMURF2* loss was predominant in EGF treated cells grown in serum containing media; in the absence of EGF, *SMURF2* knockdown caused a decrease of 21.1±2.3 and 29.3±3.4% in serum starved and serum containing conditions respectively. However, upon EGF treatment, we noted 46±3.3 and 53±4.3% decrease in EGFR surface expression depending on serum absence or presence.

As shown in **Fig. 3E**, loss of *SMURF2* resulted in significant loss of surface EGFR. In contrast, there was an increase in the cytosolic EGFR levels when the same cell was imaged using super resolution imaging on a cross sectional area of the cytosol using a greater penetration depth in the TIRF setup. However, because of clustering of EGFR, we were unable to use STORM microscopy to quantify receptor density in the cytosol. We circumvented this by investigating the relationship between fluorescence intensity of EGFR in the membrane and cytosol to that of SMURF2 in individual cells in non-reconstructed images. Two color-epi-fluorescence imaging was used to determine the fluorescence intensity of SMURF2 and EGFR in the cytosol, and TIRF imaging was used to measure the membrane fluorescence intensity of EGFR in individual cells. The intensities were then normalized and binned. As shown in **Fig. 3F**, membrane levels of EGFR are positively correlated with SMURF2 expression, whereas, cytosolic EGFR levels are inversely correlated. Together, we conclude that SMURF2 selectively protects activated EGFR.

As EGF treatment promotes endosome-mediated receptor internalization followed by lysosome-mediated degradation, we tested the effects of SMURF2 loss on EGFR cytosolic trafficking. While TIRF imaging can be used to quantify EGFR membrane expression, clustering of receptors in cytosolic organelles is less reliable. Therefore, we relied on conventional two-color colocalization studies of EGFR with EEA1 (as endosomal marker) and LAMP1 (as lysosomal marker). Pearson correlation coefficient was calculated drawing a mask of the cell and the two (control vs *SMURF2* siRNA) conditions were compared for at least 10 cells. Upon *SMURF2* knockdown, we found an increase in cytosolic EGFR colocalization both with EEA1 (*p*=0.03) and LAMP1 (*p*=0.01) (**Fig. 3G, H, and Supporting Fig. 3A, B**).

### *SMURF2* levels dictate TKI sensitivity in lung cancer cells

To understand the importance SMURF2-mediated protection of mutant EGFR in regulating TKI sensitivity, we utilized either SMURF2 overexpression or siRNA-mediated knockdown in multiple lung adenocarcinoma cell line’s sensitivity or resistance to TKI respectively. An erlotinib-sensitive PC9 cells harboring exon 19 deletion mutation in EGFR, showed increased erlotinib tolerance (IC_50_ 15.1±1.2 nM for parental, compared to 51.1±2.5 nM upon SMURF2 overexpression) using clonogenic survival assays (**Fig. 4A**). In contrast, an erlotinib resistant NCI-H1975 cells (with L858R/T790M mutations), became more sensitive (IC_50_ 5.5±1.5 μM reduced to 3.1±0.4 μM) following siRNA-mediated loss of *SMURF2* (**Fig. 4C**). Additionally, a PC9 clone resistant to third generation TKI, AZD9291 (PC9-AR), became sensitive to the drug treatment following siRNA-mediated *SMURF2* loss (**Fig. 4E**). In all these cases SMURF2 alterations also impacted EGFR steady state levels (**Fig. 4B, D, and F**) that correlated with drug sensitivity. These data supported the critical importance of SMURF2 in maintaining mutant EGFR protein stability and further identified SMURF2 targeting as a novel approach to target mutant EGFR to overcome TKI resistance.

**Figure 4.**
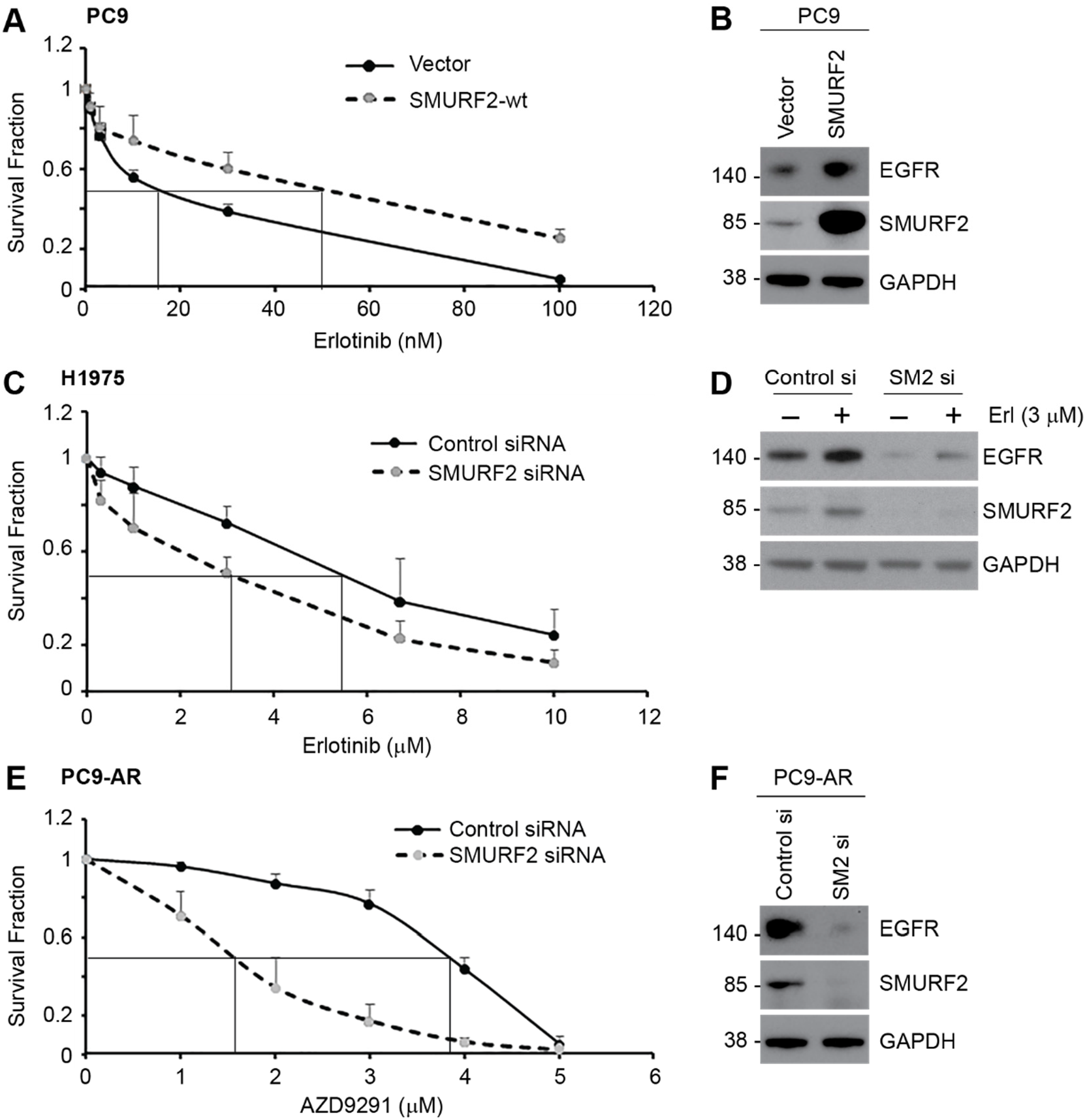
SMURF2 levels dictate TKI sensitivity in lung cancer cells. **(A)** SMURF2 was overexpressed in an erlotinib-sensitive PC9 lung adenocarcinoma cells (contain EGFR exon 19 deletion) and clonogenic cell survival assay was performed to test erlotinib sensitivity. **(B)** Immunoblot analysis of PC9 cells overexpressing SMURF2 showing increased EGFR accumulation. **(C)** Clonogenic survival of erlotinib resistant NCI-H1975 cells following control and SMURF2 siRNA-mediated knockdown. **(D)** Corresponding immunoblot analysis showing SMURF2 and corresponding EGFR loss, which correlates with the survival. **(E)** In contrast, a PC9 clone resistant to a third generation TKI (AZD9291) (PC9-AR) became sensitive to the drug treatment following siRNA-mediated SMURF2 knockdown. **(F)** A corresponding immunoblot from the above experiment confirming SMURF2 loss and its impact on EGFR.

### Alteration of SMURF2-UBCH5 protein-protein interaction impacts mutant EGFR levels

Although the finding that SMURF2 silencing can enhance mutant EGFR degradation to reduce clonogenic survival of erlotinib resistant T790-bearing lung AC cells is encouraging, direct inhibition of SMURF2 activity is not a viable strategy, due to the critical importance of SMURF2 during mitosis (35,36). Therefore, we decided to explore selective antagonism of SMURF2 interaction with its partner E2, UBCH5, to negatively impact SMURF2-mediated mutant EGFR protection from degradation. To achieve this goal, mutational analyses were carried out to define the critical residues important for SMURF2 and UBCH5 interaction. We found two (2) Proline (P) residues in UBCH5, which were critical for allowing binding between the two proteins. This is in agreement with x-ray crystal structure of SMURF2 binding with another E2 partner, UBCH7 (46). To confirm the importance of these residues, we mutated the two P residues located at 61 and 95 positions of UBCH5 to alanine (A), which surprisingly enhanced SMURF2-UBCH5 interaction compared to the wild type E2 (**Fig. 5A**). However, overexpression of such PA mutated UBCH5 caused enhanced EGFR protein degradation (**Fig. 5B**). Such studies gave us an indication as how to develop a novel strategy of targeting TKI-resistant EGFR via altering SMURF2-UBCH5 interaction.

**Figure 5.**
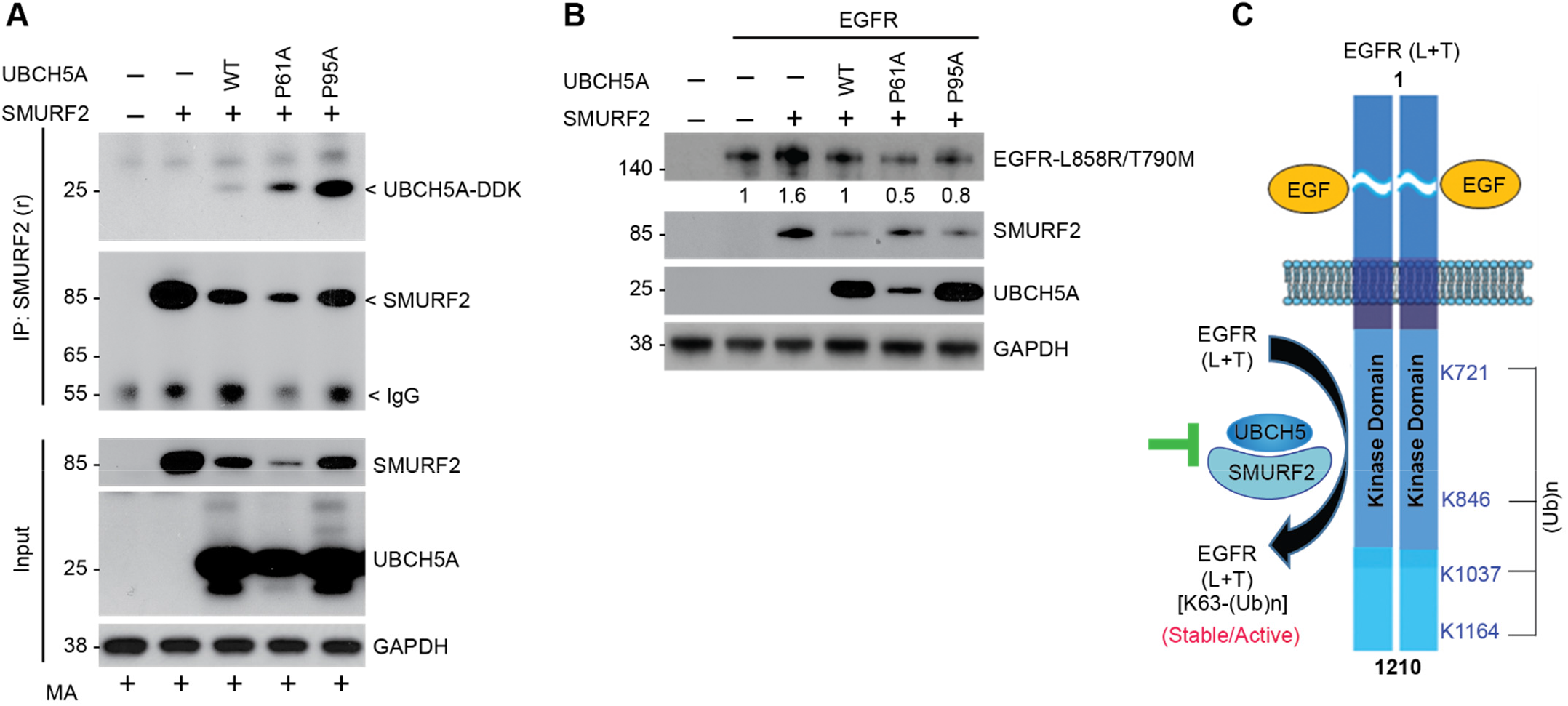
Alteration of SMURF2-UBCH5 protein-protein interaction impacts mutant EGFR levels. **(A)** Flag-tagged SMURF2 and Myc-tagged UBCH5A (wild type, P61A, and P95A mutants) were overexpressed in HEK293 cells. Cell lysates were prepared 24 h post-transfection and subjected to immunoprecipitation using SMURF2 antibody followed by immunoblotting using indicated antibodies. **(B)** Effect of overexpression of UBCH5A (wild type, P61A and P95A mutants) along with SMURF2 on L858R/T790M EGFR steady state levels. **(C)** Schematic diagram showing lysine (K) residues (K721, 846, 1037 and 1164) as identified residues underwent SMURF2-mediated ubiquitination as determined using mass spectrometry analyses. We propose altering the SMURF2-UBCH5 interaction as a future targeting strategy to promote protein destabilization of mutant EGFR to overcome TKI resistance.

## DISCUSSION

This study extends our previously reported observation that defines the critical importance of SMURF2 catalytic activity in promoting TKI resistance via mutant EGFR ubiquitination and stabilization. Here we have identified SMURF2-mediated preferential polyubiquitination of TKI-resistant (L858R/T790M) mutant EGFR as one of several mechanisms that facilitate increased membrane retention of the mutant receptor to stabilize and cause TKI resistance. Using biochemical and genetic approaches, we have identified four lysine (K) residues in T790M mutant EGFR that undergo protective ubiquitination. Mutation of one (K1037) of these lysines (a known acetylation site that increases receptor internalization) to acetylation-mimicking asparagine (Q), converted the TKI-resistant stable L858R/T790M mutant EGFR to unstable upon TKI treatment. Consequently, loss of SMURF2 enhanced mutant EGFR degradation to potentiate drug responsiveness, whereas, SMURF2 overexpression stabilized the receptor to convert a TKI-sensitive line into a resistant one. These data support the potential of SMURF2 targeting as a novel therapeutic strategy to overcome TKI resistance. We summarized such findings in **Fig. 5C**.

We previously reported that SMURF2 directly interacts with EGFR, however, unlike several other ubiquitin ligases, SMURF2’s ubiquitin ligase activity is critical for maintaining EGFR protein stability (25). Here we identified that among various mutants, L858R/T790M mutant EGFR is a preferred SMURF2 substrate, which along with its partner, UBCH5 (an ubiquitin conjugating enzyme, E2) can efficiently polyubiquitinate mutant EGFR at lysine (K) 721, 846, 1037 and K1164 positions. Conversely, siRNA mediated *SMURF2* silencing caused rapid disappearance of membrane-bound EGFR and promoted enhanced internalization and degradation (**Fig. 4, Supporting Fig. 3**). Earlier studies have indicated that K721 and K846 residues of EGFR are important for maintaining EGFR kinase activity, whereas, K1037 and K1164 are important sites of acetylation required for receptor internalization (30). In another study, Jiang et al. reported that inhibition of K1037 acetylation by an oncogenic protein, Sulfiredoxin (Srx), can promote sustained EGFR activation (37). Taken together, we hypothesize that SMURF2-UBCH5-mediated ubiquitination at K721 and K846 residues may be critical in constitutive activation of the mutant EGFR, whereas, ubiquitination at K1037 and 1164 may be competing with acetylation to balance receptor internalization, membrane retention, and protein stability. Here, we have substantiated our findings by incorporating acetylation mimicking asparagine (Q) substitution at K1037 site, which converted the erlotinib resistant L858R/T790M mutant EGFR vulnerable to degradation following drug treatment.

In this study, we further utilized both cell fractionation as well as super resolution TIRF microscopy to demonstrate the importance of SMURF2 for the activated receptor membrane retention. Our data demonstrated that via *SMURF2* targeting, we can reduce EGFR surface expression because activated receptors promptly undergo internalization and degradation via endosomal/lysosomal trafficking. While SMURF2 could be an attractive molecular target to develop anti-EGFR therapy, SMURF2 is a major mitotic regulator in spindle assembly checkpoint (SAC) (35) and forced alterations cause chromosomal instability, aneuploidy and promote enhanced tumor initiation in *Smurf2-null* mice (38). This suggests that si-/shRNA mediated silencing of SMURF2 expression may not be an ideal anticancer strategy. To increase substrate specificity without compromising SMURF2’s normal cellular function, we proposed to disrupt protein-protein interaction (PPI) between SMURF2-UBCH5. We hypothesized that as SMURF2-UBCH5 complex is specific in polyubiquitinating L858R/T790M mutant EGFR, a strategy altering the E3-E2 complex formation should impact mutant EGFR ubiquitination and promote mutant EGFR degradation. Our data revealed that there are two conserved proline (P) residues located at 61 and 95 positions of UBCH5, which are critical for SMURF2 interaction (**Fig. 5**). Mutation of either of the P to alanine (A) significantly tightened SMURF2-UBCH5 interaction and overexpression of such UBCH5 PA mutants compromised SMURF2-mediated L858R/T790M EGFR stabilization. It is known that E3-E2 interaction is dynamic and often show moderate to weak interaction (39); but as UBCH5 PA mutants bound tighter with the SMURF2 compared to wild type E2, and caused mutant EGFR degradation, we hypothesize that such tighter interaction possibly compromised the ubiquitin transfer, leading to the loss of SMURF2 mediated protective ubiquitination and degradation of the mutant receptor, similar to that found in *SMURF2* knockdown cells. Further mechanistic studies may better elucidate these interactions. Previously, Levin et al. (40) reported that mutations of F62A and A96D in UBCH5 (which are adjacent to P residues we mutated), reduced E2 catalytic activity when tested with SspH2, a bacterial E3 from *S. typhimurium*, supporting our hypothesis.

Multiple ubiquitin ligases have been implicated in maintaining EGFR protein stability including c-CBL and p-VHL (22,23). Although either siRNA-mediated loss of SMURF2, blocking of SMURF2 interaction with its cognate E2 (UBCH5), or mutation of K1037 ubiquitination site in EGFR promoted EGFR degradation upon TKI treatment, in near future it may be important to identify ligase(s) involved in L858R/T790M mutant EGFR degradation following SMURF2 targeting. Our previous studies identified SMURF2 as a negative regulator of beta-transducin repeat containing protein 1 (ß-TrCP1), an F-box family ubiquitin ligase (41). Although ß-TrCP1 has yet been implicated in promoting EGFR ubiquitination and degradation, it may be involved in targeting mutant EGFR, particularly in the absence of SMURF2. Our study also identified K721 (the catalytic lysine) and K846 in EGFR as other sites of ubiquitination, however, the importance of this ubiquitination in EGFR stability is not yet understood.

We recognize certain limitations of our study. One challenge to quantitative super resolution imaging stems from the fact that many probes used in these studies either over-count or under-count the number of labeled proteins. Labeled proteins can be under-counted if a large number of genetically encoded fluorophores fail to activate because they are improperly folded, or if proteins of interest are inaccessible to labeling antibodies. Over-counting can occur when photo-switchable or photo-activatable probes blink reversibly, or when multiple fluorophores decorate labeling antibodies. A pair-correlation based analytical method was recently developed that can correct in some instances quantitative artifacts that arise from overcounting. This method takes advantage of the observation that the magnitude of self-clustering that arises from overcounting is inversely proportional to the density of the labeled protein, as long as the labeled protein is sampled randomly. In this work, we take advantage of over-counting in super-resolution images to provide a quantitative measure of receptor density in intact cells, which we further validated utilizing conventional cell fractionation studies followed by immunoprecipitation and immunoblotting, which improved the reliability of data.

In summary, this study improved our biochemical and molecular understanding of TKI-resistance and establish SMURF2-UBCH5 mediated post-translational modifications of L858R/T790M mutant EGFR as a critical partner in TKI resistance. Furthermore, our project has the potential to develop a novel strategy capable of overcoming TKI resistance via selectively altering the interaction between SMURF2 and UBCH5 and degrading mutant EGFR.

## MATERIALS AND METHODS

### Reagents

Anti-EGFR (sc-03) and anti-ubiquitin (P4D1) antibodies were from Santa Cruz Biotechnology (Santa Cruz, CA). GAPDH antibody was purchased from Cell Signaling (Danvers, MA), whereas another EGFR antibody (31G7) and Lipofectamine were purchased from Invitrogen (Grand Island, NY). Cycloheximide (CHX) was obtained from Sigma-Aldrich (St Louis, MO), and erlotinib was obtained from Genentech Inc. (San Francisco, CA). A monoclonal antibody against human SMURF2 has been described previously (42). Rabbit polyclonal SMURF2 antibody was purchased from Sigma-Millipore. Anti UBCH5 (UBE2D1, Cat#ab66600) antibody was obtained from Abcam (Cambridge, MA). Control (Cat#D-001810) and *SMURF2* (Cat#D-007194) small interfering RNA (siRNA) were purchased from Thermo Fisher Scientific (Lafayette, CO), whereas, another *SMURF2* siRNA pool (Cat#sc-41675), were obtained from Santa Cruz Biotechnology (Santa Cruz, CA).

### Cell Cultures

EGFR-null CHO cells were purchased from the American Type Culture Collection (ATCC). The human lung adenocarcinoma NCI-H1975 was kindly provided by Dr. J. A. Engelman (Massachusetts General Hospital, Boston). Cells were cultured in RPMI-1640 supplemented with 10% cosmic calf serum (CCS). For plasmid transfection Lipofectamine and for the siRNA transfection Lipofectamine RNAimax (Invitrogen, NY, USA) were used according to the manufacturer’s instructions.

### Protein Analyses

Immunoblotting and immunoprecipitation techniques were performed as described previously (25). Briefly, cell lysates were prepared using lysis buffer containing 50 mM HEPES-KOH (pH 7.5), 150 mM NaCl, 1.3 mM CaCl_2_, 1 mM dithiothreitol, 10 mM β-glycerophosphate, 1 mM NaF, 0.1 mM sodium orthovanadate, 10% glycerol, 1% NP40 and 1X protease inhibitor cocktail (Sigma, Cat. P8340). For subcellular fractionation studies, cytosolic, nuclear, and membrane fractions were isolated using a Compartment Protein Extraction Kit (Millipore, Billerica, MA) following the manufacturer’s instructions. The extracts from these fractions were subjected to immunoprecipitation (IP), and the interaction between EGFR and SMURF2 was assessed by immunoblot analysis.

### Clonogenic cell survival assay

Clonogenic survival assays were performed using techniques described previously (43). For this assay, cells were plated at a predetermined plating efficiency and after 7-9 days colonies formed were fixed using acetic acid/methanol (1:7) followed by staining using crystal violet (0.5% w/v) solution. The effects of *SMURF2* siRNA, treatment with either erlotinib or AZD9291 treatments on clonogenic survival of different cell lines were determined by normalizing the survival fraction of control siRNA or vehicle-treated group as 1.

### *In vitro* ubiquitination assay

The *in vitro* ubiquitination reaction was carried out as described previously (41). Briefly, in reaction buffer (250 mM Tris-HCl, pH 7.5, 50 mM MgCl_2_, 50 μM DTT, 20 mM ATP), 10 μg of Myc-tagged ubiquitin (Cat#U-115), 0.35 μg of UBE1 (Cat#E305), 0.5 μg of UBCH5 (Cat#E2-616) (all from Boston Biochemicals, Cambridge, MA) were added. The human recombinant SMURF2 protein (Cat. 468H, Creative Biomart, New York, NY) and different EGFR (WT, L858R, and L858R/T790M) were then added, and the reaction mixtures were incubated at 37°C for 2 hours. The reactions were then immunoprecipitated using the EGFR antibody and the pulled immunocomplexes were boiled with 4X gel loading dye. The samples were then resolved and immunoblotted using indicated antibodies.

### Mass spectrometry

The *in vitro* ubiquitination reaction mix was separated on a polyacrylamide gel, and proteins were visualized with colloidal Coomassie stain. In-gel digestion followed by identification of ubiquitination site mapping was carried out essentially as described previously (41). Briefly, upon trypsin digestion, peptides were resolved on a nanocapillary reverse phase column and subjected to high-resolution, linear ion-trap mass spectrometer (LTQ Orbitrap XL, Thermo Fisher). The full MS scan was collected in Orbitrap (resolution 30,000@400 m/z), and data-dependent MS/MS spectra on the nine most intense ions from each full MS scan were acquired. Proteins and peptides were identified by searching the data against the UniProt human protein database (20353 entries; reviewed only) using Proteome Discoverer (v 2.4, ThermoScientific) using the following search parameters. MS1 and MS2 tolerance were set to 50 ppm and 0.6 Da, respectively; carbamidomethylation of cysteines (57.02146 Da) was considered static modification; oxidation of methionine (15.9949 Da), deamidation of asparagine and glutamine (0.98401 Da) and diglycine remnant on lysines (114.0292 Da) were considered variable. Identified proteins and peptides were filtered to retain only those that passed ≤1% FDR threshold. Spectral matches to ubiquitinated peptides were manually verified.

### Labeling

For the super resolution microscopy, cells grown in Matek glass-bottom plates. Following all treatments, cells were fixed for 10 min in PBS solution containing 4% paraformaldehyde and 0.1% glutaraldehyde. After fixing, the cells are incubated in a blocking buffer (PBS, 2% Fish Gelatin, 0.01% Sodium Azide) for one hour. The cells were then incubated in the blocking buffer containing the corresponding primary antibodies for one hour. The plates were then washed 3-5 times and then incubated in secondary antibody (Alexa 532 and 647, 1:1000) for one hour. The plates were then washed again three times with the blocking buffer.

### STORM imaging and reconstruction

Samples were imaged in one of two photoconvertible buffers. βME buffer (50 mM Tris, 0.1 g/ml glucose, 100 mM NaCl, 40 μg/ml catalase, 500 μg/ml glucose oxidase, 1% βME pH 8.5) was used for one color while 0.1M Cysteamine was used instead of βME in case of two-color STORM. A detailed description of the imaging setup and STORM reconstruction is described elsewhere (44). Briefly, dSTORM was performed on 100X UAPO TIRF objective (NA = 1.49) in an Olympus IX81-XDC inverted microscope equipped with a cellTIRF module. 532nm laser (Samba 532-150 CW, Cobolt, San Jose, CA) and a 640nm laser (CUBE 640-74FP; Coherent) were used for excitation of the two fluorophores. Images were captured on an iXon-897 Andor EMCCD camera. For two-color experiments, the emitted light was passed through a DV2 emission splitting system (Photometrics, Tuscon, AZ) to simultaneously image both the near- and far-red channels. To prevent z-drift, an active Z-drift correction (ZDC) (Olympus America, Center Valley, PA) was used. Superresolution images were reconstructed after filtering, localization of individual fluorescent events, correction of stage-drift and post-processing using our in-house MATLAB program.

### Measuring receptor density

To calculate receptor density, the reconstructed STORM image was used. Autocorrelation g(r) and crosscorrelations functions were calculated as previously described (45). At radii on the order of the resolution limit of the measurement, the autocorrelation function, Pair autocorrelation functions have a correlation, g(r)>1. In two dimensions and in the absence of significant self-clustering, the integrated intensity of the auto-correlation function is inversely related to the average surface density (ρ) of proteins accessible to labeling antibodies according to the relation:

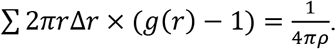

The sum is over all radius, where Δr is typically the size of the pixel in a reconstructed image and *p* is the density of receptor. Thus, larger the magnitude of autocorrelation lowers the density of receptor. The characteristic radii of the autocorrelation are related to the clustering of the receptor on the surface. To measure interaction of SMURF2 and EGFR, cross-correlation was measured for the reconstructed two-color images. The magnitude of the cross-correlation is a measure of the colocalization of the two proteins.

## Colocalization

Jacop plugin in Macphotonics ImageJ (ver 1.48) was used to measure Pearson correlation coefficient.

## Statistics

Unless noted otherwise, results are presented as mean ± SEM (standard error of mean estimate) of at least three independent experiments.

## DATA AVAILABILITY

Data described in the manuscript are located as a part of the main text, figures and the supporting figures. The mass spectrometry proteomics data have been deposited to the ProteomeXchange Consortium via the PRIDE (46) partner repository with the dataset identifier PXD018324.

## AUTHOR’S CONTRIBUTIONS

PR, KR, AA, USA, SS, VB and DR were involved in execution of works and data collection. SV, TSL, MKN and DR were involved in conceptualization, design of work, data analyses/interpretation and manuscript preparation.

## ACKNOWLEDGMENTS

This work is supported in part by grants from the National Institutes of Health R01CA160981 (to D. R.) and R01GM110052 (to S.V.). R.K. post-doctoral fellowship is funded by M-Cube, and the Fast-Forward Innovation Program at the University of Michigan. We also thank Dr. Christine Lovly from the Vanderbilt University for providing AR-resistant PC9 cells.

---

**Supporting Figure 1.**
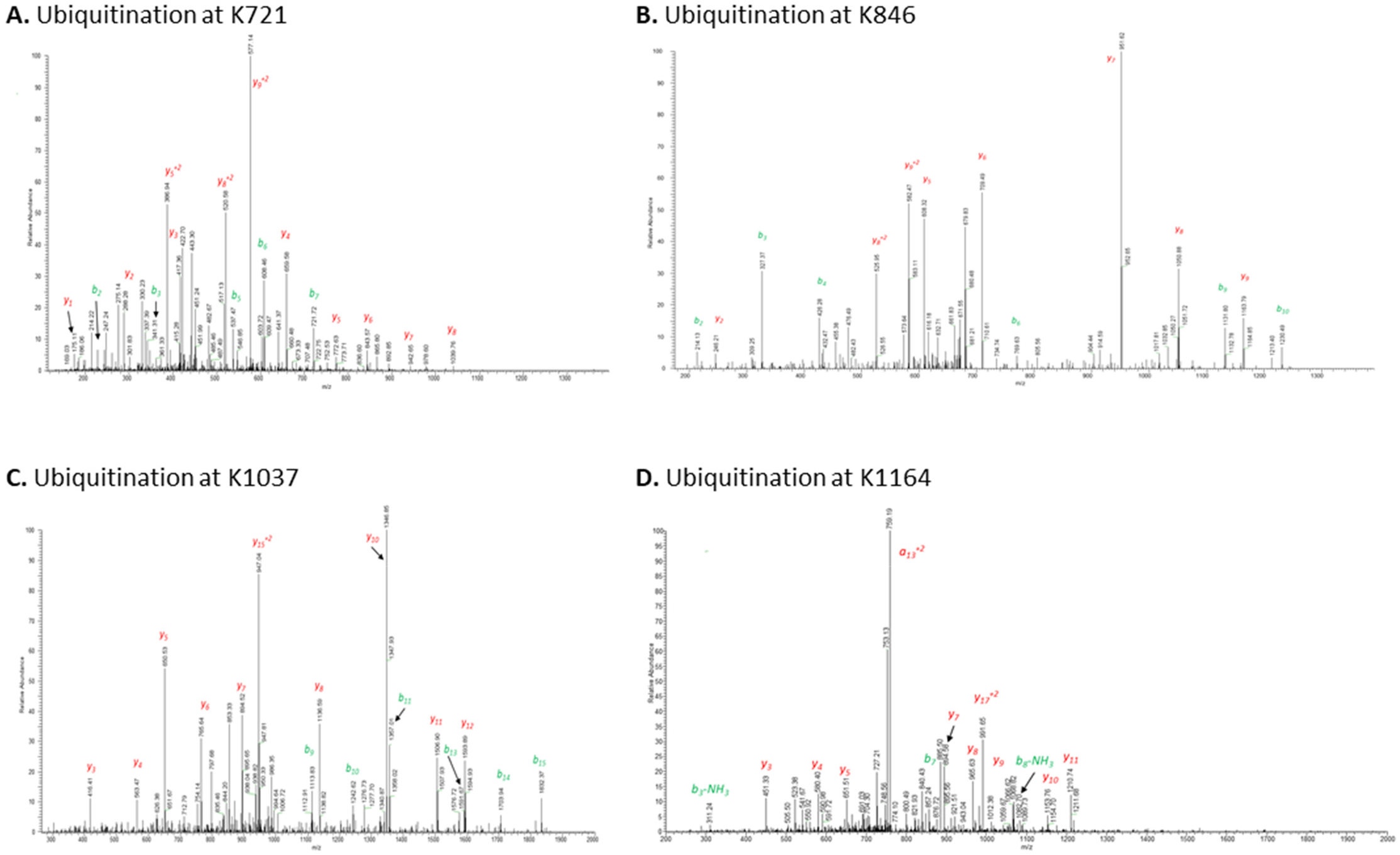
MS/MS spectrum of a peptide identifying ubiquitination of at K721, K846, K1037, K1164 sites in L+T mutant EGFR. Peptides isolated upon in-gel digestion were resolved on a reverse phase column, and collision induced dissociation spectra were obtained using an Orbitrap XL mass spectrometer. MS/MS corresponding to ^714^VKIPVAIK*ELR^724^ (in **A)** ^842^NVLVK*TPQHVK^852^ (in **B)**, ^1029^NGLQSCcamPIK*EDSFLQR^1044^ (in **C)**, and ^1156^EAKPNGIFK*GSTAENAEYLR^1075^ (in **D)** of L858R/T790M (L+T) EGFR are shown above. Observed b- and y-ions are indicated. Modified Lys (K) is denoted with *. Ccam = Carboxymethylated Cys.

**Supporting Figure 2.**
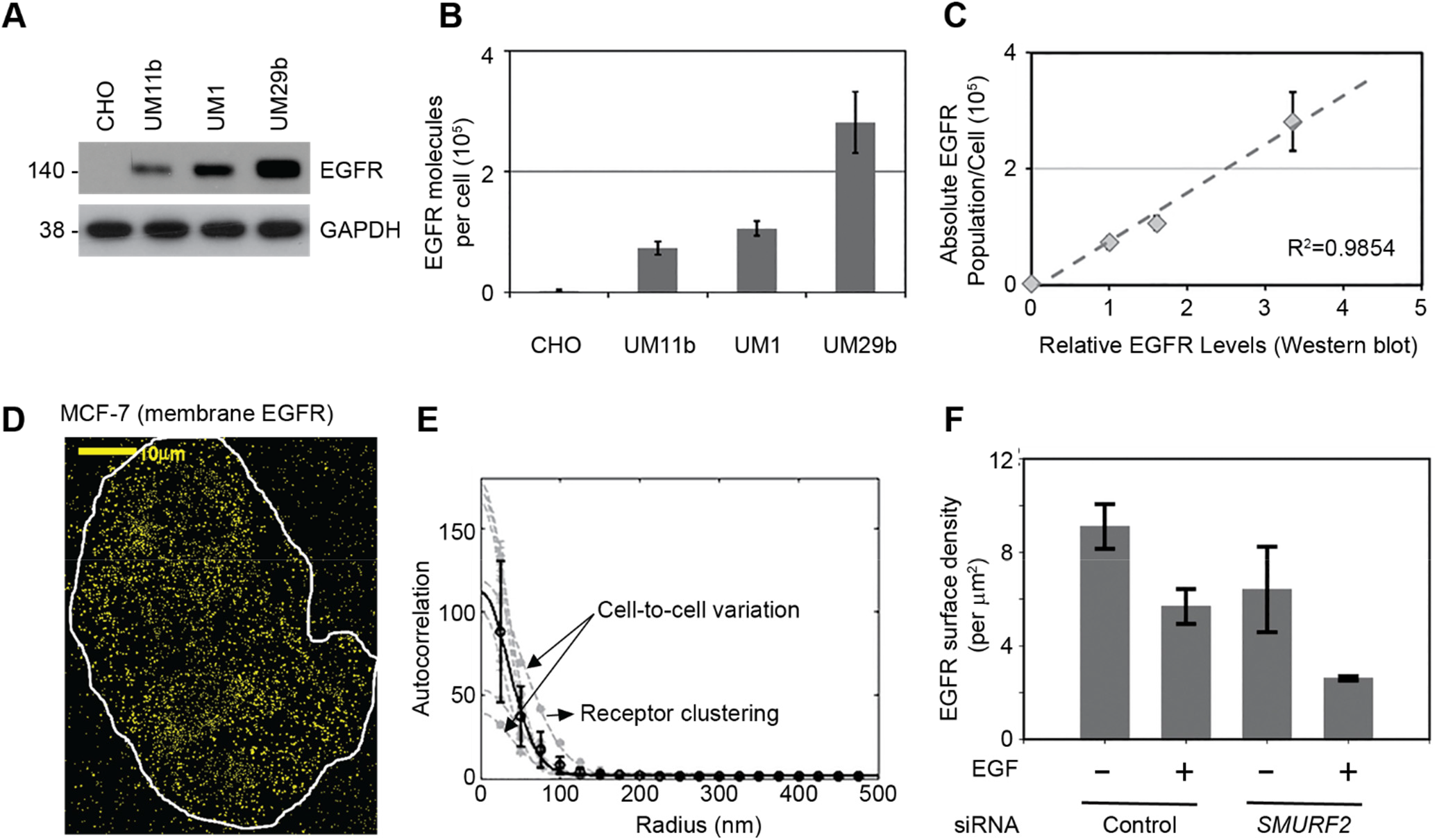
Validation of EGFR quantification in the membrane using super-resolution imaging. **(A)** Quantification of EGFR in CHO, UMSCC-11b, UMSCC-1 and UMSCC-29b cells using immunoblot analysis. **(B)** Measurements of EGFR surface population in above mentioned cell lines using super-resolution imaging. Density measured through super-resolution image is multiplied with surface area of the cell to give population per cell. **(C)** EGFR quantification using two methodologies showing strong correlation (R^2^=0.9854). **(D)** Representative reconstructed TIRF image of MCF-7 cells stained for EGFR using super-resolution imaging. Scale bar, 10 μm. **(E)** Auto-correlation functions were tabulated from images of 12 MCF-7 cells and results are summarized showing reasons of variation arise due to cell-to-cell variation and receptor clustering. **(F)** Quantification of EGFR membrane density in UMSCC-1 cells using TIRFM showing effects of EGF treatment in the presence and absence of SMURF2.

**Supporting Figure 3.**
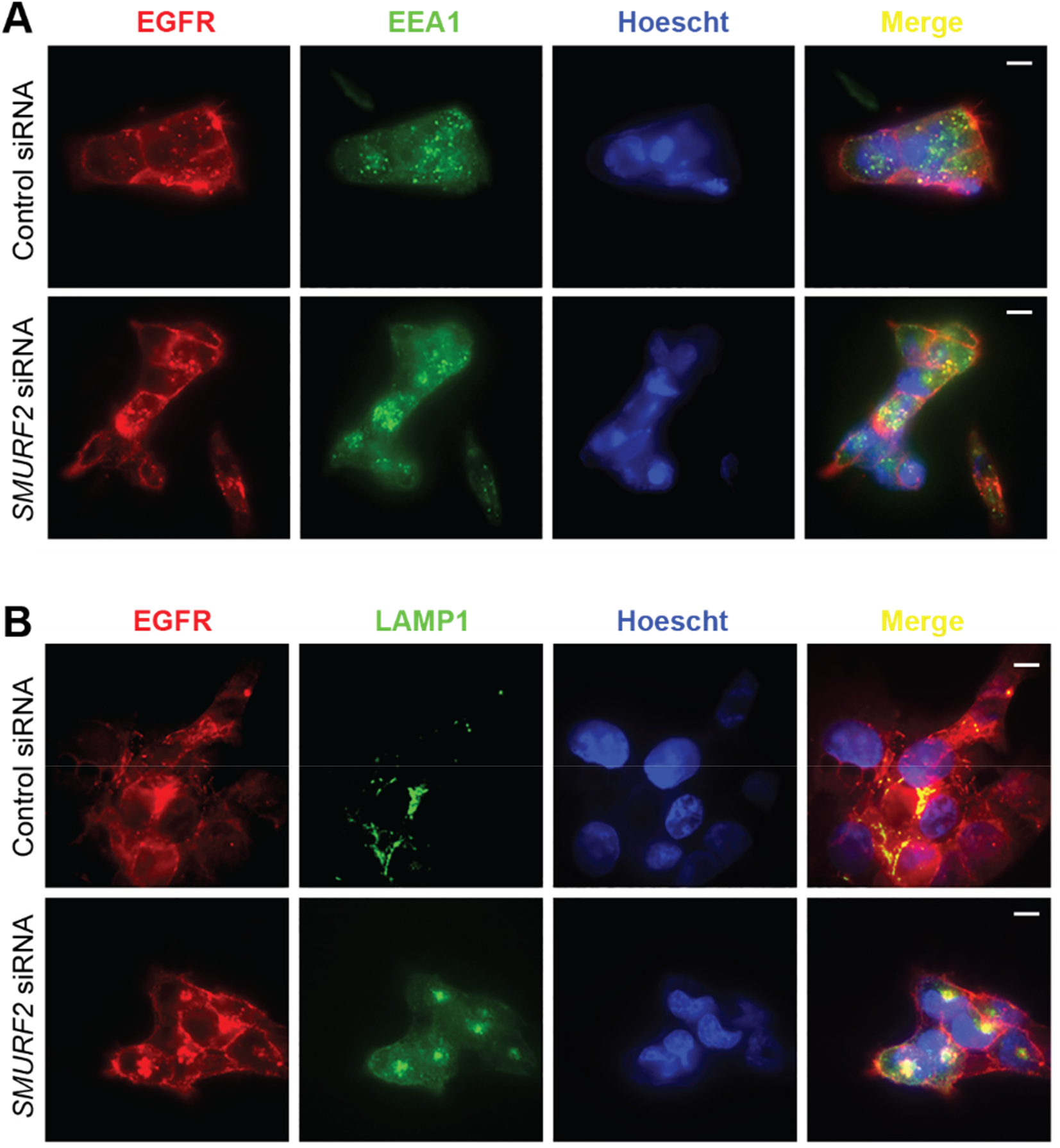
Increased localization of EGFR in the endosomes and lysosomes following SMURF2 loss. UMSCC-1 cells were either transfected with control or SMURF2 siRNA. 48 hours following transfection, cells were fixed and stained either with EGFR and EEA1 (in **panel A)**, or EGFR and LAMP1 **(panel B)** antibodies. Representative merged images are showing colocalization. Scale bar, 10 μm.

